# Sex-dependent pathways in hepatocarcinogenesis triggered by deregulated cholesterol synthesis

**DOI:** 10.1101/2020.09.03.280974

**Authors:** Kaja Blagotinšek Cokan, Žiga Urlep, Gregor Lorbek, Madlen Matz-Soja, Cene Skubic, Martina Perše, Jera Jeruc, Peter Juvan, Tadeja Režen, Damjana Rozman

**Affiliations:** Centre for Functional Genomics and Bio-Chips, Institute of Biochemistry, Faculty of Medicine, University of Ljubljana, Ljubljana, Slovenia; Rudol-Schönheimer-Institute of Biochemistry, Divison of General Biochemistry, Faculty of Medicine, University of Leipzig, Leipzig, Germany; Medical Experimental Centre, Institute of Pathology, Faculty of Medicine, University of Ljubljana, Ljubljana, Slovenia; Institute of Pathology, Faculty of Medicine, University of Ljubljana, Ljubljana, Slovenia

**Keywords:** Cholesterol biosynthesis, hepatocellular carcinoma, lanosterol 14α-demethylase (CYP51A1), sex dimorphism

## Abstract

We uncover novel pathways of sex-dependent hepatocarcinogenesis due to chronic repression of cholesterol synthesis at the lanosterol 14α-demethylase (CYP51) step. The response to metabolic insult determined by global liver transcriptome, qPCR and sterol metabolite analysis, together with blood parameters, revealed molecular signatures that differ between females and males. The data deduced from the mouse model are highly relevant for humans. The dampened hepatic metabolism presents a hallmark of carcinogenesis, particularly in ageing females, with increased plasma cholesterol and HDL, and a substantial negative enrichment of transcription factors from lipid metabolism, such as NR1B1, LXRα, LRH1, and FXR. Importantly, the carcinogenic signalling pathways (ECM-receptor interaction and PI3K/Akt) are positively enriched, albeit with sex-dependent gene targets. The activated TGF-β, mTOR, Wnt, and estrogen signalling worsen the phenotype, with NFATC1/2 being central to the female phenotype. This collectively leads to activated cell death and diminished basal metabolism. In conclusion, our data underline sex as an important biological variable of hepatocarcinogenesis. We uncover novel cholesterol-dependent transcription factors and signalling pathways as cancer markers in the ageing females.

**Significance:** Chronic repression of the late part of cholesterol synthesis provokes hepatocarcinogenesis with sex-dependent modulation of signalling pathways and transcription factors. Aging females show specific metabolic signatures and a more aggrevated phenotype of metabolism-related HCC.

## INTRODUCTION

Metabolic abnormalities result in metabolic associated fatty liver disease (MAFLD), previously termed (NAFLD) that affects over 30% of the western population and has no approved drug therapies (1). MAFLD can further develop into hepatocellular carcinoma (HCC) and both diseases are predicted to increase in the following decades (2) not only in men but also in women (3).

Among lipid factors associated with progressive liver damage is also cholesterol. The cellular cholesterol concentrations are fine-tuned by a balance of synthesis, uptake, and efflux (4). Genes of cholesterol synthesis differ in their susceptibility to loss of function mutations and can affect cell metabolism in different manners, from promoting cell survival towards manifesting in malignancies (5). Linking cholesterol synthesis and liver disease remains controversial. Cholesterol synthesis was shown to support the growth of HCC lesions after depletion of fatty acid synthesis (6), indicating a cross-talk between fatty acid and cholesterol pathways in carcinogenesis. Overexpression of the squalene epoxidase enzyme of cholesterol synthesis was identified as a driver of NAFLD-induced HCC (7). On the other hand, blocking cholesterol synthesis at the lanosterol 14α-demethylase (CYP51) step leads to the prepubertal onset of liver injury with ductular reaction and fibrosis and a pronounced male prevalent liver dysfunction before the puberty (8). In young adults, the liver damage was more prominent among female mice due to diminished cholesterol esters and elevated sterol substrates (9).

The sex disparity in liver pathologies is far from conclusive. While some studies don’t describe sex as a cofounding factor (6, 7, 10, 11), there is increasing evidence that after the menopause MAFLD occurs at a higher rate in women compared to men (12). Female and male livers are metabolically distinct organs, which was demonstrated also in mathematical models of the liver with unique regulators of sex-specific metabolic outcomes (13). A recent review concluded that clinical and epidemiological studies frequently fail to address sex differences appropriately (12).

This study aimed to address the long-term effect of disrupted liver cholesterol synthesis where the knock-out model of ageing mice was applied. Surprisingly, the tumors developed spontaneously mainly in female ageing mice. We identified novel metabolic pathways and transcription factors that can be linked to metabolism associated with female-prone hepatocarcinogenesis and provide mechanistic insights with relevance for humans.

## RESULTS

### (1) Long-term hepatocyte deletion of Cyp51 results in HCC

We monitored the liver pathology in mice between 12 month and 24 month of age. Moderate to severe ductular reaction bridging the adjacent portal tracts was observed, accompanied by mild inflammation (predominantly mononuclear cells; arrows in **Fig. 1A**), and fibrosis that was more pronounced in females (Sirius red staining (SR); **Fig. 1B**). Yellow pigment likely representing lipofuscin (arrow in **Suppl. Fig. 1**) was observed in areas of pronounced ductular reaction, sometimes surrounded by other hepatic cells, evaluated as macrophages (stars in **Fig. 1A**).

**Figure 1.**
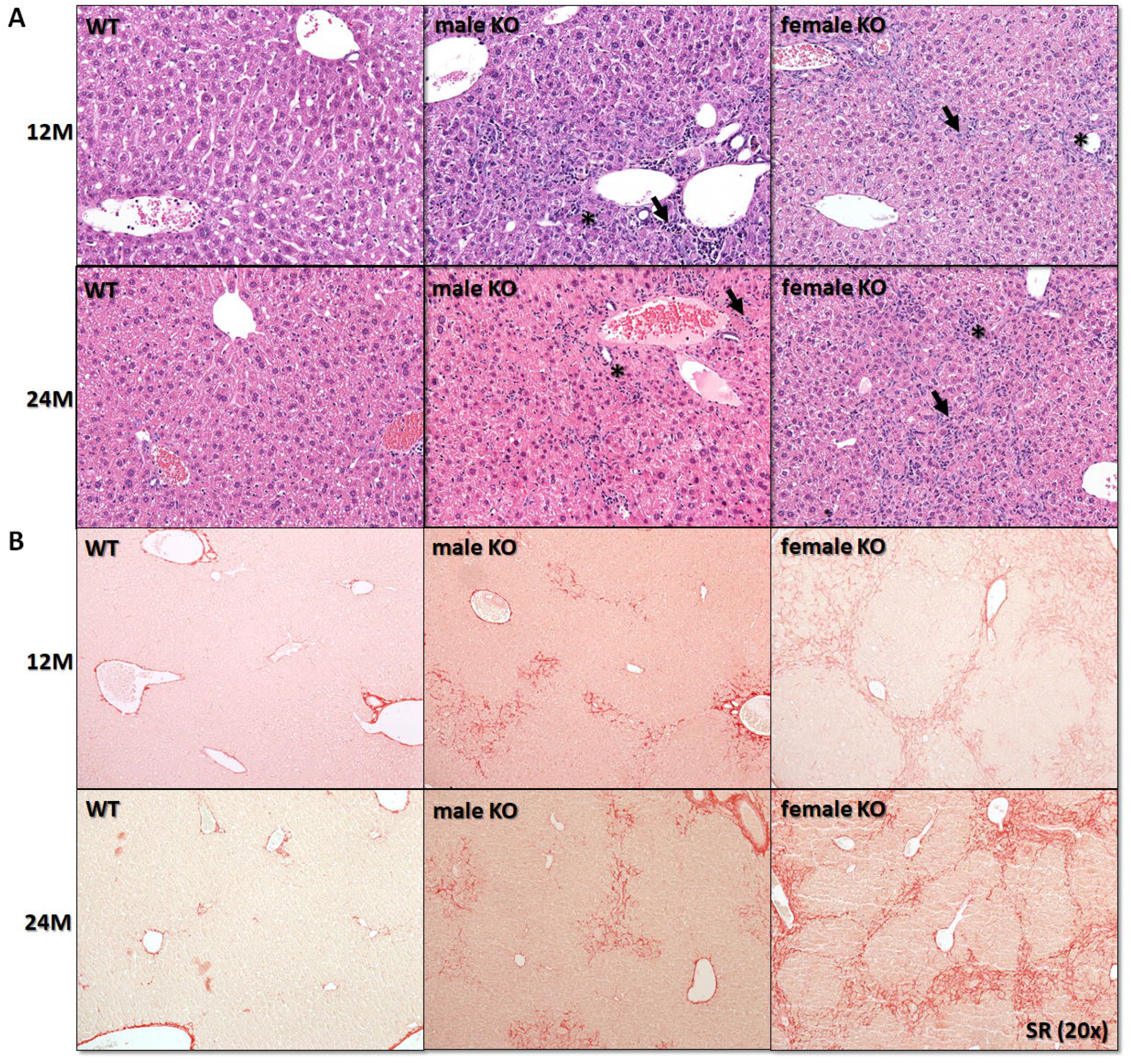
Blocking cholesterol synthesis in KOs resulted in histological alterations of the liver, particularly among females. (**A**) Histological examination revealed ductular reaction (**\***) and inflammation (→) (H&E, original magnification x200), N=5-10. (**B**) Sirius red stain (SR, original magnification x200) showed more pronounced fibrosis in female transgenic mice, N=3-5. M, month; KO, *Cyp51* KO; WT, *Cyp51* WT.

Surprisingly, liver tumors were observed in 12M female and 18M male *Cyp51* KO mice (**Fig. 2A**) defined as macroscopically visible nodules (> 1 mm) of various colours. At 24M the incidence of tumors was 77.8% for KO females and 50% for males. In WTs, a single tumor was found in a 2-year male, histologically described as adenoma (**Suppl. Table 2**). At 24M, the KOs showed a decreased body weight compared to controls, significantly higher liver weight, and elevated relative liver weights especially in females with the liver tumors (**Suppl. Fig. 2 & Suppl. Table 3**). Tumors were histologically classified as eosinophilic/clear cell nodules, hepatocellular adenomas, or HCCs (15). Immunohistochemical staining for glutamine synthetase as a recently proposed marker of HCC is shown in **Suppl. Fig. 3**. For further molecular analyses we selected only tumors histologically classified as HCC, with features such as the absence of portal tracts and hepatic lobules, thickened hepatic plates, solid growth, focal tubular or pseudoglandular structures, focal necrosis, numerous apoptotic cells, and presence of cellular and nuclear polymorphism with brisk mitotic activity (**Fig. 2B, C**).

**Figure 2.**
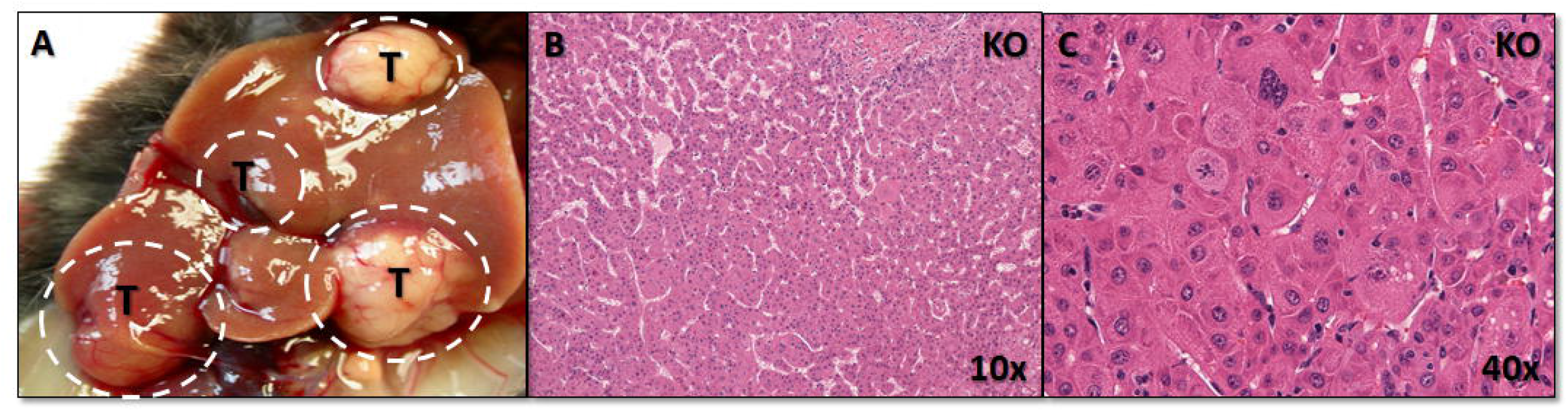
Blocking cholesterol synthesis in KOs resulted in hepatocarcinogenesis, the bigger tumor histologically consistent with HCCs. (**A**) Representative gross image of liver tumors in 24M KO mice. (**B**) Representative histologic image of HCC showing different architectural abnormalities. **(C)** Higher magnification showing cellular and nuclear polymorphism with brisk mitotic activity. Hematoxylin and Eosin (H&E), original magnification **A, B** x100, **C** x400. KO, *Cyp51* KO; T, tumor.

**Figure 3.**
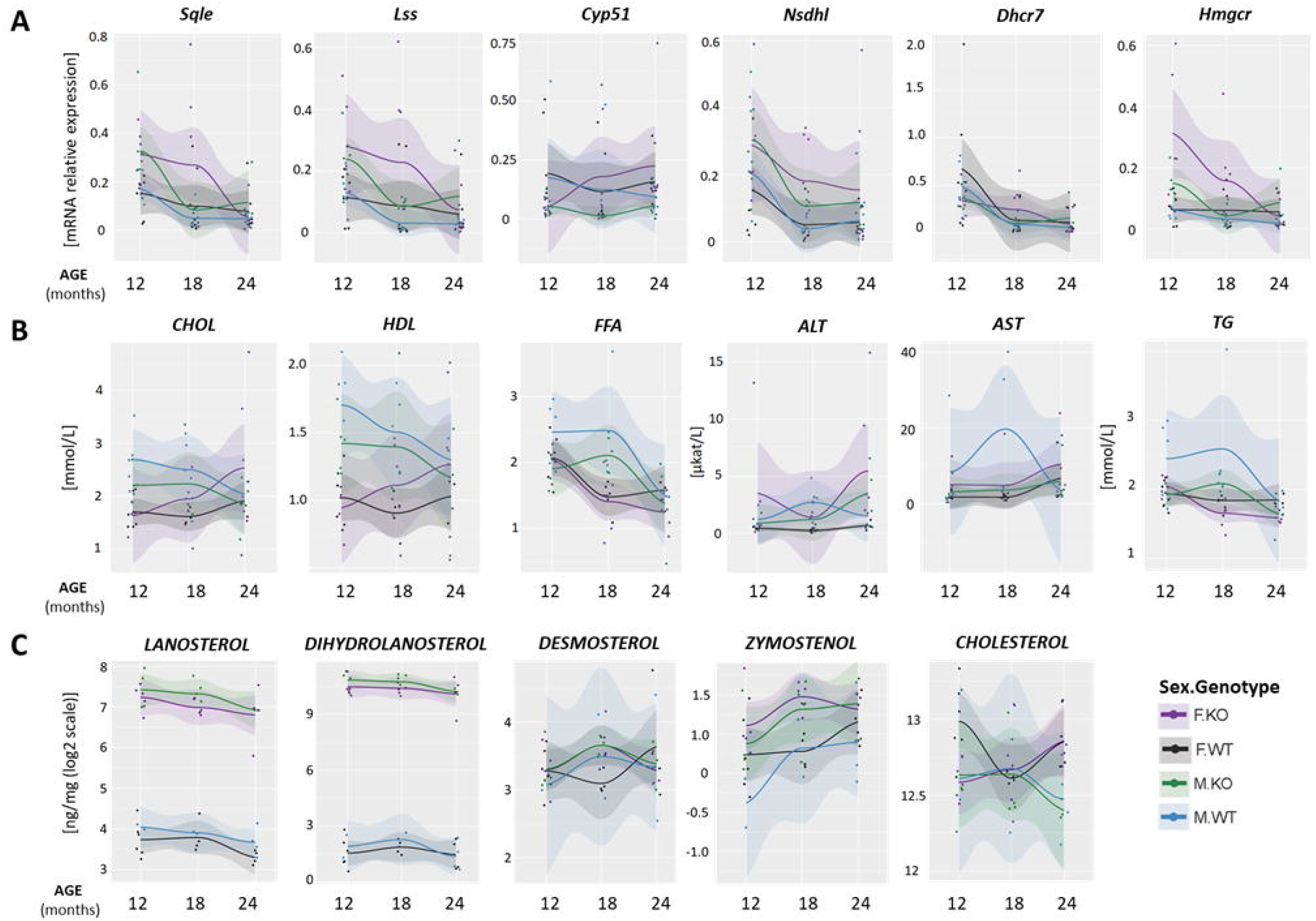
Age profiles of the compensatory changes in cholesterol homeostasis in KOs and WTs. (A) Expression of hepatic genes from cholesterol synthesis N = 5 mice per sex/genotype/age group. (**B**) Plasma biochemical parameters, N=5-10 mice per sex/genotype/age group. (**C**) Levels of selected hepatic sterol intermediates and cholesterol, N=4-6 mice per sex/genotype/age group. Light color bands - 95% confidence interval. Dots - individual measurements. KO, *Cyp51* KO; WT, *Cyp51* WT, F = female; M = male.

### (2) From metabolism associated features of MALD towards HCC in females and males

The long-term ablation of *Cyp51* in hepatocytes was studied at the gene and metabolic levels. Deregulation of genes from cholesterol synthesis was more pronounced in the *Cyp51* KO females (**Fig. 3A & Suppl. Tables 4 - 6**). Additionally, expression of *Cyp8b1, Cyp7a1* and *Cyp27a1* from the bile acid (BA) synthesis pathway was at 24 months also decreased only in female KO mice, indicating a drop in the classical pathway of BA synthesis (**Suppl. Fig. 4 & Suppl. Tables 4 - 6**).

**Figure 4.**
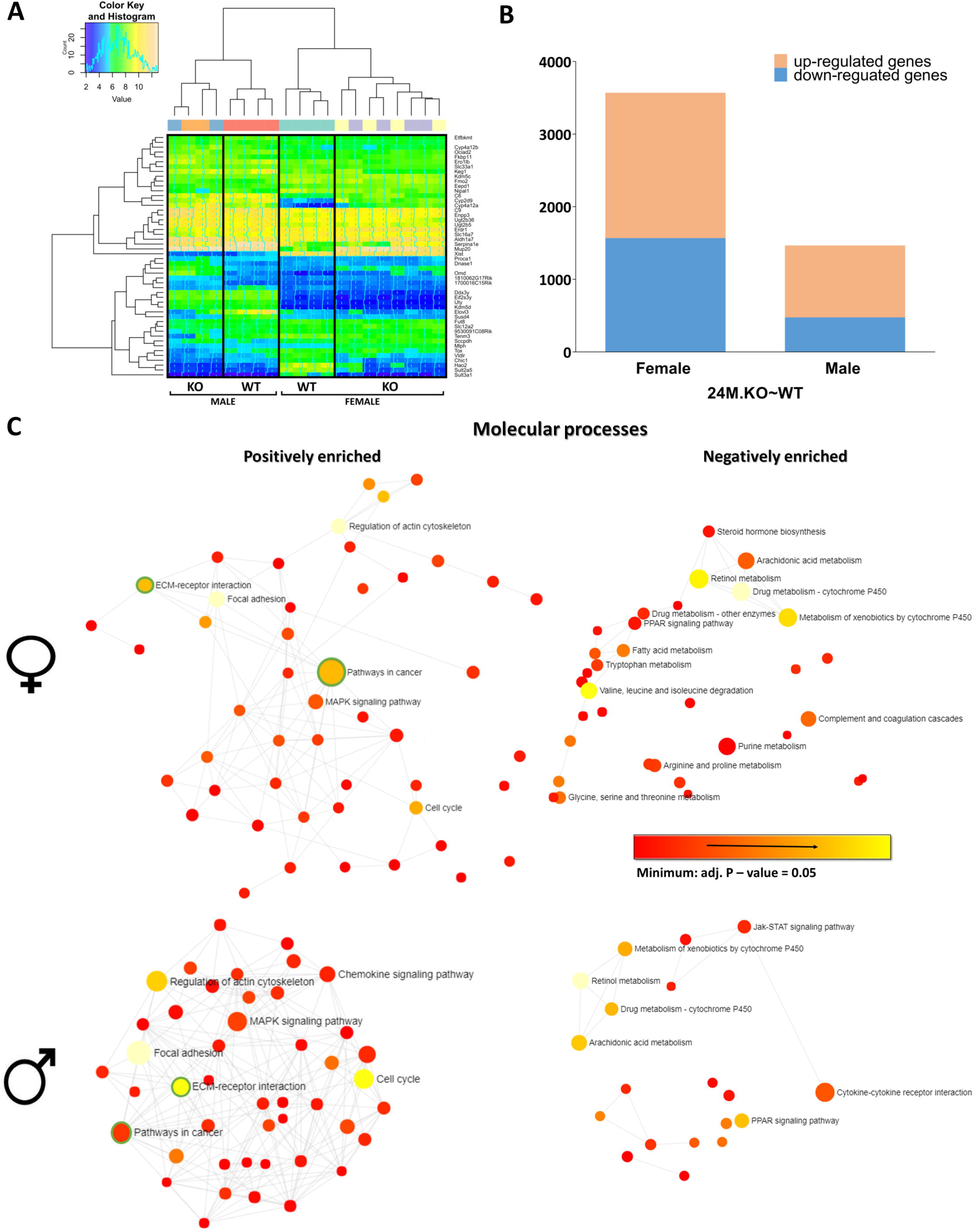
Target genes associated with blocking cholesterol synthesis. (**A**) Heat-map of top 50 DE transcripts that showed significant differences in at least one comparison: hierarchical clustering distinguished between the genotypes and sex of 24M.KO∼WT mouse livers. (**B**) Stacked column plots with numbers of differentially up- and down-regulated sex-specific genes of 24M *Cyp51* KO mice. (**C**) The functional enriched networks of up- and down-regulated DE genes in female and male 24M.KO∼WT mice using NetworkAnalyst. The size of nodes corresponds to the number of DE genes within a particular molecular process. All pathway nodules are differentially expressed (p<0.05), the gradation rising from red to yellow (most enriched) in the network of positively (left) and negatively (right) enriched molecular processes. Circles representing cancer pathways and ECM-receptor interaction are surrounded by green lines.

Plasma cholesterol (CHOL), free fatty acids (FFA), HDL (high-density lipoprotein), and triglycerides (TG) showed opposite trends among females and males during the ageing, with CHOL and HDL increasing in KO females. A significant increase in ALT in female *Cyp51* KO mice is an additional indication of chronic hepatic injury (**Fig. 3B** & **Suppl. Table 4**).

In the livers, we measured CHOL and the intermediate sterols of the cholesterol synthesis (**Fig. 3C** & **Suppl. Table 4**). In the KO mice with inactivated CYP51A1 enzyme, its substrates lanosterol and dihydrolanosterol were highly elevated (∼100-fold and ∼500-fold, respectively), proving disrupted cholesterol synthesis. Dihydrolanosterol was higher than lanosterol indicating that 24-dehydrocholesterol reductase (DHCR24) enzyme functions normally. Downstream the pathway, desmosterol was not statistically significantly different between *Cyp51* KO and WT mice, however, zymostenol was significantly higher in *Cyp51* KO livers. Small or no change between genotypes indicates that sterol intermediates were replaced by transport to the liver or by the upregulated synthesis in other cell types.

For liver CHOL, sex represents a statistically significant parameter in the interaction analysis between sex and age. Similarly as for blood CHOL also hepatic CHOL is increased in female mice during the ageing while this was not the case for the males **(Fig 3B, C & Suppl. Table 4)**.

### (3) Transcriptome analysis identified sex differences in gene expression linked to hepatocarcinogenesis

We used Affymetrix microarrays for gene expression profiling of 24M *Cyp51* KO (24M.KO) and controls (24M.WT) mouse liver samples. Data was compared to 19W *Cyp51* KO (19W.KO) and controls (19W.WT) (14) (**Suppl. Tables 6 – 8**) and discussed in the view of the early processes towards carcinogenesis.

Hierarchical clustering of differentially expressed (DE) genes showed a clear difference between 24M.WT and 24M.KO mice for both sexes (**Fig. 4A**) and a clear sex imbalance in the number of DE genes (**Fig. 4B**). From these, 11.6% DE genes including *Trp53, Ezh2, Foxa1*, and *Rb1* were deregulated only in males, while 63.4% DE genes including *Ctnnb1, Gsk3b, Nova1, Lmna, Gstp1*, and *Mup20* were deregulated only in females (**Suppl. Tables 6, 9**). Furthermore, a majority of DE genes were positively enriched (56% in females and 68% in males) (**Fig. 4B**). Using NetworkAnalyst (16), the functionally enriched networks of DE genes showed that positively enriched genes were mostly involved in pathways related to cancer and extracellular matrix (ECM)-receptor interaction pathway, while genes from metabolic pathways were negatively enriched, especially among female KOs (**Fig. 4C**).

### (4) Enriched KEGG pathways and transcription factors (TFs) as triggers of female prevalent hepatocarcinogenesis

From 323 KEGG pathways, we discovered 209 that were enriched in livers of 19W.KO mice aligned with 19W.WT (**Suppl. Tables 7, 10**) and 230 in 24M.KO compared to 24M.WT mice (**Table 1, Suppl. Table 7**), including the activation of HCC, apoptosis, endocytosis, PI3K/Akt signalling, pathways in cancer, and proteoglycans in cancer. ECM-receptor interaction pathway was observed as the most positively enriched in 19W KO females (**Suppl. Table 10**) and 24M KO females and males (**Table 1**), which is in line with the histological findings (**Fig. 2B**). Chronic injury induces numerous molecular alterations in hepatocytes and causes a gradual accumulation of extracellular matrix components (ECM) that might lead to a complete architectural reconstruction of hepatocytes and tumor development.

**Table 1.** The most significantly enriched KEGG pathways and their regulation by DE TFs in 24M KOs. Parameters shown in bold are statistically significant. Dark green logFC - the most negative enriched pathway/TF, dark violet logFC – the most positive enriched pathway/TF. KEGG, Kyoto Encyclopaedia of Genes and Genomes; * p<0.05, ** p<0.01, *** p<0.001.

| KEGG PATHWAY | 24M.KO~WT |  |  |  | TRANSCRIPTION FACTOR | 24M.KO~WT |  |  |  |
| --- | --- | --- | --- | --- | --- | --- | --- | --- | --- |
|  | FEMALES |  | MALES |  |  | FEMALES |  | MALES |  |
|  | LogFC | adj. P-value | LogFC | adj. P-value |  | LogFC | adj. P-value | LogFC | adj. P-value |
| *Cell cycle* | 1.60 | <0.001<br>*** | 2.13 | <0.001<br>*** | SP1 | 1.47 | <0.001<br>*** | 1.49 | <0.001<br>*** |
| *Pathways in cancer* | 1.89 | <0.001<br>*** | 1.84 | <0.001<br>*** | C-JUN | 1.42 | <0.001<br>*** | 1.50 | <0.001<br>*** |
| *ECM-receptor interaction* | 1.82 | <0.001<br>*** | 1.85 | <0.001<br>*** | JUND | 1.05 | <0.001<br>*** | 0.91 | <0.001<br>*** |
| *Proteoglycans in cancer* | 1.63 | <0.001<br>*** | 1.61 | <0.001<br>*** | C-FOS | 1.47 | <0.001<br>*** | 1.33 | <0.001<br>*** |
| *Apoptosis* | 1.65 | <0.001<br>*** | 1.49 | <0.001<br>*** | AP1 | 1.17 | <0.001<br>*** | 0.88 | <0.001<br>*** |
| *p53 signalling pathway* | 1.26 | <0.001<br>*** | 1.33 | <0.001<br>*** | C/EBPβ | 0.91 | <0.001<br>*** | 0.87 | 0.004<br>** |
| *MicroRNAs in cancer* | 0.87 | <0.001<br>*** | 0.95 | <0.001<br>*** | Egr1 | 0.95 | <0.001<br>*** | 1.03 | <0.001<br>*** |
| *Hepatocellular carcinoma* | 0.95 | <0.001<br>*** | 0.72 | 0.006<br>** | SOX9 | 0.90 | <0.001<br>*** | 0.63 | 0.001<br>** |
| *Circadian entrainment* | 0.42 | 0.020<br>* | 0.27 | 0.197 | (SOX9)2 | 0.67 | 0.005<br>** | 0.50 | 0.062 |
| *TGF-β pathway* | 0.45 | 0.007<br>** | 0.14 | 0.427 | HIF1α | 0.43 | 0.008<br>** | 0.58 | 0.004<br>** |
| *Wnt signalling pathway* | 0.32 | 0.018<br>* | 0.24 | 0.118 | NFATC1 | 0.36 | 0.035<br>* | 0.29 | 0.156 |
| *PI3K/Akt signalling pathway* | 1.61 | <0.001<br>*** | 1.51 | <0.001<br>*** | CLOCK | -0.26 | 0.031<br>* | -0.41 | 0.009<br>** |
| *Fatty acid degradation* | -1.18 | 0.001<br>** | -1.12 | 0.009<br>** | LXRα:RXRα | -0.56 | 0.043<br>* | -0.19 | 0.602 |
| *Metabolic pathways* | -1.80 | 0.009<br>** | -0.69 | 0.367 | HNF4α | -1.60 | <0.001<br>*** | -1.17 | 0.019<br>* |
| *Peroxisome* | -1.82 | <0.001<br>*** | -1.20 | 0.018<br>* | RORα |  |  |  |  |
| *Primary bile acid biosynthesis* | -1.29 | 0.002<br>** | -1.40 | 0.004<br>** | RORC | -1.68 | <0.001<br>*** | -2.10 | <0.001<br>*** |
| *Biosynthesis of unsaturated fatty acid* | -0.87 | 0.018<br>* | -0.46 | 0.261 | NR1B1 | -0.80 | <0.001<br>*** | -0.59 | <0.001<br>*** |

Interestingly, the TGF-β signalling pathway was significantly elevated only in *Cyp51* KO female livers (19W: **Suppl. Table 10**; 24M: **Table 1**). Additionally, the Wnt signalling and circadian entrainment pathways were activated only in KO females (**Table 1**). We further explored if these pathways could serve as triggers for female hepatocarcinogenesis in the *Cyp51* KO model. The selected protein markers of the TGF-β and Wnt signalling were evaluated by qPCR and IHC. Here TGF-β showed a positive reaction, particularly in female *Cyp51* KO mice (**Suppl. Fig. 5, 6** & **Suppl. Table 11**). Biosynthesis of unsaturated fatty acids, fatty acid degradation, and peroxisome pathways were negatively enriched in KO females at 19W and 24M as well as males at 24M KO (19W: **Suppl. Table 10**; 24M: **Table 1**).

**Figure 5.**
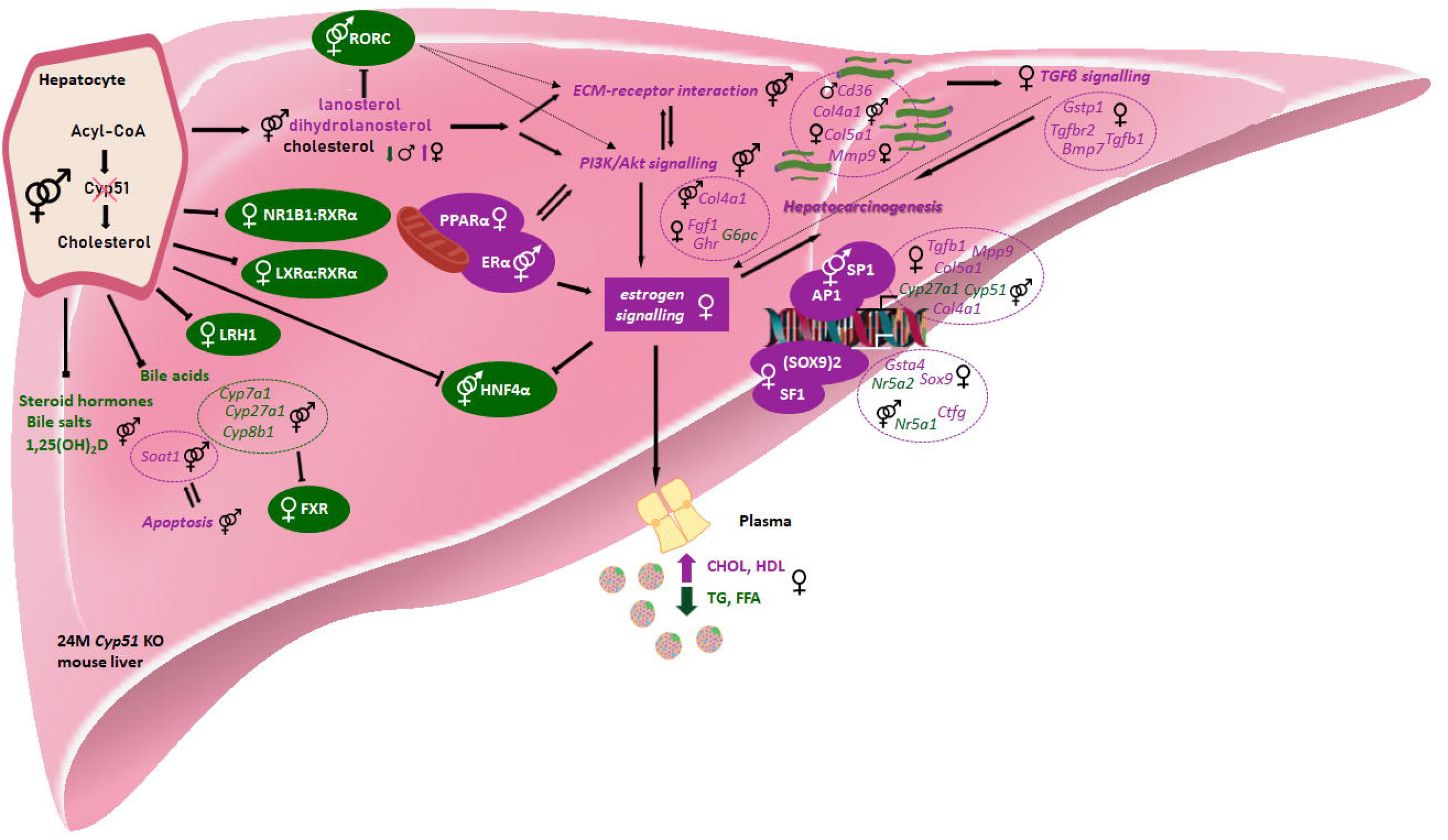
Sex-dependent metabolic and transcriptional changes after disrupted *Cyp51* from cholesterol synthesis align with hepatocarcinogenesis. CYP51A1 enzyme was repressed in hepatocytes of both sexes. However, chronic metabolic insult resulted in a variety of sex-specific metabolic changes. In the liver of both sexes at 24 months, we observed accumulation of hepatic lanosterol and dihydrolanosterol (metabolites of CYP51A1 enzyme), while hepatic cholesterol is increased in females and decreased in males, both WTs and KOs. There is a substantial negatively enrichment of TFs related to metabolism in female *Cyp51* KO mice, such as the NR1B1:RXRD, LXRD:RXR, LRH1, and FXR. The carcinogenesis related signalling pathways (ECM-receptor interaction and PI3K/Akt) are positively enriched, with sex-specific gene targets. The activated TGF-β and estrogen signalling pathways worsen the phenotype, particularly among females. Increased plasma CHOL and HDL, and decreased plasma TGs and FFA, are observed only in female KO mice. Arrows indicate active connections between enriched genes, KEGG pathways, and TFs. Blocked arrows represent repression. Hatched arrows represent possible transcription regulation of the selected pathway or connection between selected pathways in KO mice. Metabolites, DE genes, KEGG pathways, and TFs are violet if positively enriched and green if negatively enriched. Icons were used from the Reactome icon library. All selected mouse signalling and metabolic pathways and TFs were aligned with human literature data.

Considering the general alteration of DE genes and KEGG pathways in *Cyp51* KO mice, the transcription landscape was investigated. The TF enrichment analysis indicated 321 and 142 enriched TFs in 19W.KO and 24M.KO in comparison to WT littermates (**Suppl. Table 8**). The enriched TFs with specific functions are listed in **Suppl. Table 10** (19W), and **Table 1** (24M). In both groups, the transcriptional activity of RORC was negatively enriched (at 19W.KO in females only), while SOX9, SP1, AP1, and C-JUN TFs activities were positively enriched. The oncogenic isoform of SOX9 ((Sox9)2) was positively enriched only among females. The expression of the majority of RORC and SOX9 target genes was decreased in *Cyp51* KO mice, while *Cd36* in *Cyp51* KO 19W males and 24M of both sexes was positively enriched (**Suppl. Table 12**). Additionally, the RORC gene target *G6pc*, which is involved in the regulation of the PI3K/Akt signalling pathway, was significantly negatively enriched in female KO mice (**Suppl. Table 12**).

The TF networks were visualized with STRING (17) to determine the interactions. Different colours of nodes represent biological processes in which enriched TFs of *Cyp51* KO mice are involved. Surprisingly, SP1 and JUND expressed in both sexes represent nodules with most protein-protein associations linked to hepatocarcinogenesis of *Cyp51* KO mice (**Suppl. Fig. 7**).

## DISCUSSION

The hallmark of cancer cells is metabolic reprogramming, causing new cellular demands in selective survival and growth (18). Cholesterol is a precursor of steroid hormones and bile acids and an essential component of cellular membranes. Its circulating levels are under healthy homeostatic conditions regulated by a balance between cellular cholesterol synthesis, dietary intake, and removal of excess cholesterol (19).

*CYP51A1* is a gene from the late (post-squalene) part of the pathway (20). The disruption of this gene in hepatocytes starts with morphological alterations, such as ductular reaction accompanied by mild inflammation, and can end with malignant liver tumors in ageing mice that are first observed at 12M in females and 18M in males, with the highest incidence in the 24M female *Cyp51* KO mice. Also, humans show sex-dependent differences in liver pathologies. Women more commonly present with acute liver failure, benign liver lesions, autoimmune hepatitis, and toxin-mediated hepatotoxicity. Oppositely, malignant liver tumors, and viral hepatitis are less commonly observed in women (21). However, non-alcoholic steatohepatitis (NASH) as a more serious form of MALD, more commonly progresses to HCC in women (22). Generally, women are less prone to developing HCC than men (due to hormone-dependent resistance), while after the menopause a sharp increase in HCC incidence is witnessed (23).

The latest studies indicate that post-squalene intermediates of cholesterol synthesis are also signalling molecules. Inhibiting different enzymes from the pathway results in the accumulation of sterol intermediates with different cellular effects. Sterol products can activate RORC signalling (24), promote oligodendrocyte formation (25), or activate LXRα and inhibit EGFR signalling (26). Inhibitors of lanosterol synthase were identified as potential anticancer drug targets (27) while the accumulation of lanosterol can cause degradation of 3-hydroxy-3-methyl-glutaryl-coenzyme A reductase (HMGCR) (28). Our sterol measurements show significant accumulation of CYP51A1 substrates lanosterol and dihydrolanosterol, with differences between males and females regarding lanosterol. The main difference between the genotypes is likely in the accumulation of sterol substrates rather than a lack of downstream sterols, while the difference between the sexes is shown by the sex-dependent transcriptome response to sterol imbalance. Due to the lack of sterols between lanosterol and zymosterol as proposed RORC natural ligands, we observe at 24M the repression of the RORC signalling pathway in both sexes. The RORC-dependent mechanisms in cancerogenesis were recently described. *Oh et al*. reported that *RORC* expression is decreased in basal-like subtype cancers and inversely correlated with histological grade and cancer drivers in breast cancer cohorts (29). In line with our findings, negatively enriched RORC could inversely regulate the TGF-β/EMT signalling in *Cyp51* KO mice.

Injured hepatocytes in *Cyp51* KO mice produce different ECMs, exemplified by enrichment of laminin and collagen genes (*Col5a1, Col4a1, Lamc1*, etc.) and strongly positively enriched ECM-receptor interaction pathway in KOs of both sexes. In some way, these alterations could contribute to microenvironmental reorganization and induction of epithelial-mesenchymal transition (EMT) that positively influences tumor progression (30). A higher rate of cancer progression in female *Cyp51* KO mice could be explained by the female-specific interplay of positively enriched TGF-β signalling. An activated TGF-β represents a central regulator of chronic liver disease and could contribute to the progression of all disease stages in animal models and humans (31). Interestingly, important crosstalk between TGF-β and *Cyp51* gene expression has been shown in the *Tgf-*β*fl/fl*; Wnt-Cre mouse model (32).

Furthermore, the results of our analysis identified the key transcriptional regulators of DE molecular pathways: positively enriched SOX9, SP1, and negatively enriched RORC. SOX9 is an important candidate for the cancer marker required for the activation of the TGF-β/Smad signalling (33). As SOX9, the tumor progression driver SP1 is also a key factor for the induction of TGF-β during inflammation (34) and its overexpression in *Cyp51* KO relates to EMT and migration of pancreatic cancer cells (35). The impact of higher disease progression in female *Cyp51* KO mice is reflected also in the interplay of transcription regulator SMAD3 on TGF-β signalling (36).

Increased plasma CHOL and HDL, and decreased plasma TG and FFA, are observed only in female KO mice. This is a possible effect of induced sex hormone-binding globulin (SHBG) by specific targets of an estrogen signalling pathway (37). SHBG is positively correlated with HDL and negatively with TGs and has been already associated with different cancers in ageing humans (38, 39). Supporting other hepatocarcinogenesis studies (40, 41), the outcome of altered cholesterol and BA homeostasis was a chronic liver injury in our *Cyp51* KO mouse model with widespread fibrosis, inflammation, and malignant transformation due to toxic sterol accumulation. Additionally, sterol accumulation, in our case the consequence of inhibition of CYP51A1 (14), could be a cause of the observed widespread increase of hepatic progenitors (also known as a ductular reaction) that may also differentiate into mature hepatocyte-like cells with oncogenic potential (42).

Based on our data we propose that the repression of CYP51A1 enzyme can contribute to HCC development. In humans, whole-body deletion of *CYP51A1* is not reported, since this gene is essential and is embryonically lethal in the mouse (43). Only two studies are reporting the *CYP51A1* deleterious mutations, which resulted in infant liver failure and death (44, 45). Numerous polymorphisms in *CYP51A1* have been associated with different phenotypes in humans (46), but the consequences of CYP51A1 heterozygosity in human hepatocarcinogenesis need yet to be established. *CYP51A1* has a low mutation rate, lower than other cholesterol synthesis genes (47). The *CYP51A1 rs37417517* variant was found associated with the female-specific breast cancer and variant *rs229188* with the women ageing genetics (48, 49). Additionally, the COSMIC database revealed the presence of somatic *CYP51A1* point mutations in tumor samples, some also with predicted deleterious effect on activity (**Suppl. Tables 13, 14**).

## CONCLUSIONS

The hallmark of cancer cells is metabolic reprogramming, causing new cellular demands in selective survival and growth. Innactivation of *CYP51A1* from cholesterol synthesis provokes global transcriptome changes. Multiple signalling pathways and transcription factors are deregulated in a sex-dependent manner (**Suppl. Fig. 8)** which leads towards metabolic-related HCC. Female-specific cholesterol-related mechanism of hepatocarcinogenesis was aligned with human data (**Fig. 5**) and is in line with the fact that the liver is a sex-dimorphic metabolic organ and a major organ of cholesterol homeostasis. A novel finding of our work is that cholesterol synthesis deregulation due to repression of *Cyp51* initially results in NASH-like features that progress towards hepatocellular carcinoma in a sex-specific manner, with more aggrevated phenotype and specific signalling pathways in the aging females.

## MATERIALS AND METHODS

Detailed materials and methods are described in **Supplementary Materials and methods**. The experimental design of the study is shown in the **Suppl. Fig. M1**.

### Animals

The homozygous hepatocyte-specific *Cyp51* knock-out (*Cyp51* KO) and wild-type (*Cyp51* WT) mice of both sexes were used following relevant guidelines and regulations. The mice were sacrificed with cervical dislocation at 12, 18, and 24 months (M). Organs were removed, patho-morphology examined, freshly frozen, and stored at – 80 °C.

### Ethics statement

Mouse experiments were approved by the Veterinary Administration of the Republic of Slovenia (license numbers 34401-31/2011/4 and 34401-52/2012/3) and were conducted in agreement with the National Institute of Health guidelines for work with laboratory animals, national legislation and Directive 2010/63/EU on the protection of animals used for scientific purposes.

### Histology

Hematoxylin and eosin (H&E) staining, Sirius red (SR) staining and immunohistochemical reaction for glutamine synthetase, TGF-β and β-catenin were carried out on 4-5 µm formalin-fixed and paraffin-embedded liver sections.

### Analysis of plasma parameters

The concentration of lipids (total and HDL cholesterol, FFA, TG) and activity of liver enzymes (ALT and AST) were measured in plasma by commercial kits.

### Liver sterol analysis

Sterol intermediates were isolated as described (50) and identified and quantified by LC-MS. After normalization based on internal standard, the concentration of individual sterols was calculated corresponding to the standard sterols from Avanti Lipids.

**Gene expression profiling** of liver samples of 24M old *Cyp51* WT and KO mice of both sexes was performed using Affymetrix GeneChip(tm) Mouse Gene 2.0 ST arrays and validated by qPCR. Samples of 20 mice were obtained from normal livers as controls (wild-type, WT) and macroscopically visible liver tumors with surrounding tissue as KO. The data were analysed using R/limma controlling false discovery rate at α=0.05 and deposited to GEO under accession number GSE127772. KEGG PATHWAY and TRANSFAC databases were used for functional enrichment studies. Gene expression data from 19 weeks (W) (14) WT and KO mice of both sexes were used to determine processes that contribute to tumor progression from 19W towards 24M. Network diagrams were created using NetworkAnalyst and STRING. The proposed mechanistic scheme was designed from data obtained from mice experiments and aligned to human literature data to increase their relevance.

## DECLARATIONS

### Ethical Approval and Consent to participate

None.

### Consent for publication

None.

### Availability of supporting data

The datasets generated during the current study are available in the Gene Expression Omnibus database (https://www.ncbi.nlm.nih.gov/geo/) under accession number GSE127772 and GSE58271.

### Author contributions

Study design and supervision: D.R.; Experiment conduct: K.B.C., G.L., M.P., C.S.; Data analysis: P.J.; Data evaluation and interpretation: K.B.C., M.P., J.J., C.S, D.R.; Material support: M.M.S.; Writing of the manuscript: K.B.C., M.P., J.J., T.R., P.J., D.R. All the authors contributed to the final version of the manuscript.

## Acknowledgements

We thank K. Kodra, H. Klavžar, M. Kužnik, and D. Mahn for their help with experiments. We acknowledge S. Horvat and R. Keber for their previous contribution to the development of the *Cyp51* knock-out mouse model. Thanks also to N. Nadižar for help in statistical data evaluation and figures preparation and P. Ivanuša for help with sterol isolation. Thanks to collaborators I. Vovk and M. Križman for LC-MS method development.

