## Supplementary material for "Sex-dependent pathways in hepatocarcinogenesis triggered by deregulated cholesterol synthesis": Suppl. Material and methods. Suppl. Figures.

***SUPPLEMENTARY MATERIALS AND METHODS AND SUPPLEMENTARY FIGURES***

***Supplementary materials and methods***

**Suppl. Fig. M1:** Experimental design.

**Suppl. Table M1:** List of negatively tested microorganisms in mice.

**Suppl. Table M2:** Number of mouse samples for plasma and RT-qPCR gene

expression analysis.

**Suppl. Table M3:** Number of 24M mouse samples for DNA-microarray

analysis.

#### Suppl. Table M4: Number of 19W mouse samples from GSE58271 for DNA-microarray analysis.

***Supplementary Figures***

**Suppl. Fig. 1:** Patho-histological features of *Cyp51* KO mice.

**Suppl. Fig. 2:** The KO mice liver and body weights at different ages, both sexes.

**Suppl. Fig. 3:** Immunohistochemical expression of glutamine synthetase.

**Suppl. Fig. 4:** Relative mRNA expression of selected genes of cholesterol and	BA synthesis, the concentration of plasma biochemical parameters	and, sterol liver intermediate cholesterol.

**Suppl. Fig. 5:** Immunohistochemical expression of TGF-β1 and β-catenin*.*

**Suppl. Fig. 6:** qPCR expression profiles of TGF-β and Wnt signalling markers.

**Suppl. Fig. 7:** TF networks of 24M KO mice.

**Suppl. Fig. 8.** Hepatocyte-specific *Cyp51* knock-out resulted in the deregulation	of multiple signalling pathways and TFs leading to the development	of female-prevalent liver cancer.

***SUPPLEMENTARY TABLES***

**Suppl. Table 1:** List of gene sequences for qPCR validation.

**Suppl. Table 2**: Female *Cyp51* KO mice had a higher incidence of liver tumors as males.

**Suppl. Table 3:** Differences between mouse liver/body/relative weights.

**Suppl. Table 4:** A heat-map of cholesterol and bile acid synthesis DNA-‍microarray gene expression in male and female KO mice.

**Suppl. Table 5:** The long-term ablation of *Cyp51* in hepatocytes influence at gene and metabolic levels.

**Suppl. Tables 6:** Gene expression tables.

**Suppl. Tables 7:** KEGG pathways expression tables.

**Suppl. Tables 8:** Transcription factors expression tables.

**Suppl. Table 9:** A clear sex imbalance in gene expression in 24M KOs.

**Suppl. Table 10:** Enriched KEGG pathways and transcription factors in male and female mice.

**Suppl. Table 11:** qPCR expression profiles of TGF-β1 and Wnt signalling markers.

**Suppl. Tables 12:** Target genes of TFs RORC and SOX9.

**Suppl. Tables 13:** COSMIC database information of *CYP51A1* point mutations in humans.

**Suppl. Tables 1:** The positive data (mutated samples) for selected gene *CYP51A1* from COSMIC database.

***SUPPLEMENTARY MATERIALS AND METHODS***

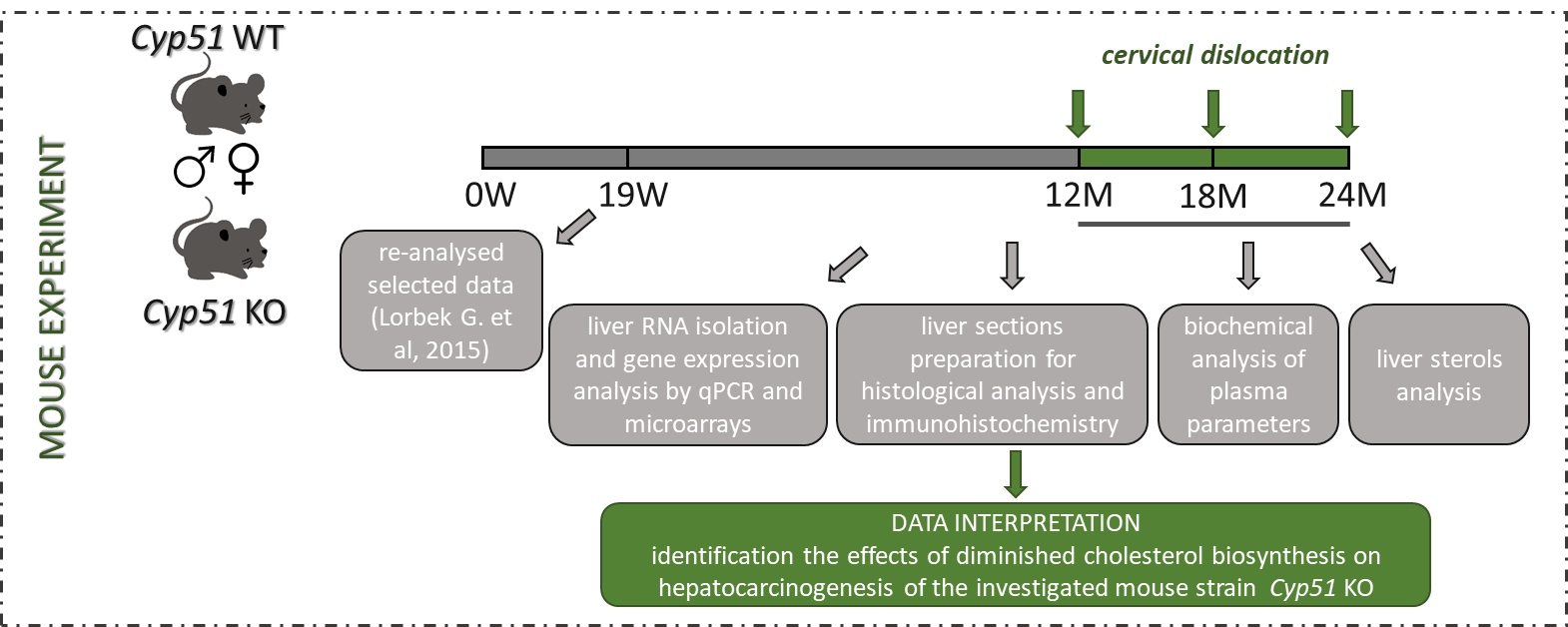

#### **Suppl. Fig M1:** Experimental design. Experiments were performed in female and male *Cyp51* liver conditional knock-out mice at age of 12, 18, and 24 months, and the wild-type controls. Collected data from livers and blood samples were aligned to data of the 19 weeks old mice.

**Animal study and samples collection.** The generation of homozygous hepatocyte-specific *Cyp51* knock-out (*Cyp51* KO) and wild-type (*Cyp51* WT) mice on a mixed genetic background (129/Pas (10%) × C57BL/6J (90%)) was established by crossbreeding between transgenic mouse strains with floxed exons 3 and 4 of the *Cyp51* allele (1) and transgenic mice expressing recombinase *Cre* under the control of the albumin promoter (Cre-Alb) as has been reported previously (2, 3). Animals were bred under controlled conditions: a 12h light and dark cycle (7:00 am till 7:00 pm light), the temperature of 22±1 ºC, humidity 55±10%, maintained on standard rodent chow (diet 1324, Altromin, Germany) and acidified tap water (pH = ‍3) *ad libitum*. The experiment was performed at the Medical Experimental Center, Faculty of Medicine, University of Ljubljana. During the experiment, the routine health microbiological monitoring was done. Mice were negative for the microorganisms stated in **Table M1**. Mice were housed in groups (3-5 mice per cage) in open bar cages (825 cm^2^ floor area, Ehret, Germany) on bedding material (lignocel ¾, Germany) enriched by paper towels and mouse houses (Tecniplast, Italy). Genotypes were confirmed by polymerase chain reaction (PCR) of ear or tail gDNA as was previously described (2).

We sacrificed *Cyp51* KO and *Cyp51* WT mice of both sexes between 11h and 13h after a 4-6h of fast (food withdrawal at 7h a.m.) with cervical dislocation at different time points of age: 12, 18 and 24 months. Immediately after cervical dislocation we took blood from the right ventricle, collected into heparin-coated Vacuette MiniCollect® 1 ml Plasma Tubes (Greiner Bioone, Frickenhausen, Germany) and centrifuged for 4° C, 3000 g for 15 min. Mice were weighed, organs (liver, kidney, spleen, heart, gonads) were removed and patho-morphologically examined. For histology, the left lateral lobus was fixed in 4% formalin and embedded in paraffin for further histological analysis. Other parts of livers were cut into thin slices and snap-‍frozen in liquid nitrogen. The material was freshly frozen and stored at – 80 ‍°C for subsequent analysis.

Suppl. Table M1: List of negatively tested microorganisms in mice in routine health microbiological monitoring.

| **List of negatively tested microorganisms** | |
| --- | --- |
| *Citrobacter rodentium* | Minute virus of mice |
| *Corynebacterium kutscheri* | Mouse hepatitis virus |
| *Klebsiella oxytoca* | Mouse parvovirus (rVP2) |
| *Klebsiella pneumoniae* | Mouse rotavirus / EDIM |
| *Pseudomonas aeruginosa* | Mycoplasma pulmonis |
| *Salmonela spp.* | Pneumonia virus of mice |
| *Staphylococcus aureus* | Reovirus type 3 |
| *Streptobacillus moniformis* | Sendai virus |
| *Streptococci beta-haemolytic Group A* | Theiler's encephalomyelitis virus (GD VII) |
| *Streptococci beta-haemolytic Group B* | Myobia musculi / Radfordia sp. |
| *Streptococci beta-haemolytic Group C* | Other ectoparasites |
| *Streptococci beta-haemolytic Group G* | Aspiculuris tetraptera |
| *Streptococcus pneumoniae* | Cryptosporidium spp. |
| *Adenovirus FL* | Entamoeba spp. |
| *Adenovirus K87* | Giardia spp. |
| *Clostridium piliforme* | Helicobacter bilis |
| *Ectromelia virus* | Spironucleus spp. |
| *General parvovirus (rNS-1)* | Syphacia obvelata |
| *Lymphocytic choriomeningitisvirus* |  |

**Histological analysis.** For standard histological analyses formalin-fixed paraffin-embedded tissue was used, sectioned to 4-5 µm and deparaffinized at 70 °C for 10 min and washed in two changes of Xylene, 100% ethanol, 95% ethanol, 70% ethanol and twice in acidified water. Hematoxylin and Eosin (H&E) stained liver sections of each mouse were scored individually for different parameters such as size, shape, and polymorphism of the hepatocytes, granulation of cytoplasm, presence of portal and parenchymal inflammation, ductular reaction, cholestasis, and presence of any type of nodules. The intensity of inflammation and ductular reaction was graded as follows: 0 - normal, + - mild, ++ - moderate and +++ - severe. The extent of ductular reaction was assigned as follows: p – proliferation around portal tracts, p-mz – proliferation extending from one portal tract to the middle zone, and p-p – bile ducts were extending from one portal field to another – bridging.

For evaluation of liver fibrosis, Sirius red (SR) stain was used. SR staining was performed by incubating deparaffinized sections in 0.1% Sirius Red solution for 1h (SR; 0.1% direct red 80, 1.2% picric acid in water), briefly destaining in diluted acetic acid, dehydrating in 70% ethanol, 95% ethanol, 100% ethanol and twice in Xylene and fixed with Roti Histokitt II (Carl Roth GmbH + Co. KG, Germany). Fibrosis in SR stained liver samples was graded as follows: 0 – absent, + mild periportal, ++ moderate, and +++ as very strong fibrosis with bridging.

Tumors, identified as macroscopically visible nodules (> 1 mm) of various colors observed at autopsy, were histologically classified based on Mouse tumor classification criteria (4) as eosinophilic or clear cells nodules, adenomas, cholangiomas, cholangiocellular carcinoma or hepatocellular carcinoma (HCC).

**Immunohistochemistry.** To evaluate the expression of selected markers in liver samples submitted to the microarray experiment, immunolabeling for glutamine synthetase, β-catenin, and TGF-β1 was performed. Liver tissue sections were deparaffinized and subjected to degrading alcohol gradient and treated for antigen retrieval before staining. Afterward, 3% hydrogen peroxide (H_2_0_2_) was used for endogenous peroxidase quenching, non-specific staining was blocked using 5% normal goat serum (Sigma-‍Aldrich, St. Louis, MO, USA) for 1h at room temperature (RT). Incubation with primary, anti-glutamine synthetase (BD Biosciences, San Jose, California, USA; 1:1000 dilution), anti-‍β-‍catenin (BD Biosciences, San Jose, California, USA; 1:1000 dilution) and anti-TGF-β1 (Promega, Madison, WI, USA; 1:500 dilution) antibodies in 1% goat serum in 0.1% TBST was performed overnight at 4 °C. We used DAKO EnVision Detection System (Agilent Technologies DAKO, Glostrup, Denmark) for antibody detection. The sections were subsequently counterstained with hematoxylin.

**Plasma analysis.** Mice were placed into groups according to their age (12, 18 and 24 months, 24 mice per group), genotype (37 *Cyp51* WT, 35 *Cyp51* KO), and sex (40 female, 32 male). 72 mouse samples (4-7 samples per group) were analysed (**Table M2**).

#### Suppl. Table M2: Number of mouse samples for plasma and RT-qPCR gene expression analysis.

| **Mouse samples** | | Age | | |
| --- | --- | --- | --- | --- |
| Genotype | Sex | 12M | 18M | 24M |
| *Cyp51* WT | female | 7 | 7 | 7 |
|  | male | 5 | 5 | 6 |
| *Cyp51* KO | female | 5 | 7 | 7 |
|  | male | 7 | 5 | 4 |

Total and HDL cholesterol, free fatty acids (FFA), triglycerides (TG), alanine aminotransferase (ALT), and aspartate aminotransferase (AST) were analyzed by Veterinary ambulance Moste (Ljubljana, Slovenia) with Architect ci8200 analyzer (Abbott Diagnostics, Abbott Park, IL, USA). The concentration of lipid parameters were measured as mmol/L and a linear regression model *Age+Genotype+Sex+Age:Sex* was used to fit the concentrations of individual lipids. The activity of ALT and AST were given in μkat/L and a linear regression model *Age+Genotype+Sex+Genotype:Sex* was used to fit their activity individually. ANOVA type III test was used to estimate the statistical significance of the effects and the Holm method was used to control the family-wise error rate (FWER) at α=0.05.

**Liver sterol analysis.** Sterol isolation: Sterol intermediates were isolated with the protocol described previously (5). Homogenized mouse livers were weighted and transferred to Folch solution (chloroform:methanol = 2:1) and incubated at room temperature for 24h. Folch solution with samples corresponding to 80 mg of liver tissue was transfer to fresh tubes and 200 ng of internal standard Lathosterol-D7 (Avanti Polar Lipids) was added. Folch was dried in a vacuum centrifuge, 1ml of Hydrolysis solution was added and shaken for 2h in a water bath at 65 ◦C. MiliQ water (0.5 ml), and cyclohexane (3 ml) were added, vortexed and centrifuged at 3500 rpm in for 10 min. The upper phase with sterols was transferred into a new 15 ml tube and the extraction was repeated once more with 3 ml of cyclohexane. Extracts were pooled and evaporated in vacuum centrifuge. Samples were dissolved in 250 µl of methanol and transferred to HPLC vials for analysis.

LC-MS analysis: Sterols intermediated were separated on two combined columns Luna® 3 µm PFP (2) 100 Å (100 mm and 150 mm length) on Shimadzu Nexera XR HPLC. For mobile phase, 80% methanol, 10% water, 10% 1-propanol, and 0.05% Formic acid were used in the isocratic condition. HPLC was coupled with the Sciex Triple Quad 3500 mass spectrometer for detection. For ionization APCI (Atmospheric-pressure chemical ionization) was used in positive mode and detection was made in MRM (Multiple reaction monitoring) mode. The sample concentration was normalized on internal standard (Lathosterol-d7) and sterol concentrations calculated to corresponding standards from Avanti Lipids. We measured following sterol intermediates with MRM in bracket: Zymosterol* (367/215), 24-dehydrolathosterol* (367/215), 7-‍dehydrodesmosterol* (365/199), Desmosterol (367/215), Zymostenol (369/215), Lathosterol-d7 (376/215), FF-MAS* (393/214), T-MAS* (395/243), Cholesterol (369/215), Lanosterol (409/191) and 24,25-dihydrolanosterol (411/191). *Sterol concentration was too low to quantify.

**Gene expression analysis.** The same mice and groups were used for plasma analysis (**Table ‍M2**). Total RNA was isolated from 30 µg of the liver using TRI Reagent (Sigma-Aldrich, St. Louis, MO, USA) procedure. RNA concentration and purity of each sample were assayed using NanoDrop 1000 Spectrophotometer (Thermo Fischer Scientific, Waltham, MA, USA). The RNA quality was checked with Agilent 2100 BioAnalyzer (Agilent Technologies, Santa Clara, CA, USA). Before cDNA synthesis, all liver RNA samples (2 µg) were treated with amplification grade DNase I (Roche, Basel, Switzerland), and reverse-transcribed with the Transcriptor Universal cDNA Master (Roche, Basel, Switzerland) according to manufacturer’s instructions.

Real-time quantitative reverse transcription polymerase chain reaction (RT-qPCR) was performed using a Roche LightCycler 480 (Roche, Basel, Switzerland). In all experiments, the PCR reaction consisted of 2.5 µL SYBR Green I Master (Roche, Basel, Switzerland), 0.75 µL of cDNA template, 0.6 µL of 2.5 µM random primer mix and 1.15 µL of RNAse-free water in a final volume of 5 ‍µL. All experiments were carried out in three technical replicates for each of the sample using 384-well plates. PCR was performed with the following parameters: 10 ‍min incubation at 95 °C followed by 45 cycles of 10 s at 95 °C, 20 s at 58 °C and 20 s at 72 ‍°C. *Utp6* and *Hmbs* were chosen as internal reference genes in mice liver for normalization of gene expression data using NormFinder (6). The relative expression ratio was calculated using the ddCp as previously described (7). The list of primer sequences used in RT-‍qPCR is provided under **Suppl. Table 1**: oligonucleotide sequences. A linear regression model *Age+Genotype+Sex+Age:Genotype* was used to fit normalized expression data of cholesterol-‍related genes *Hmgcr, Sqle, Lss, Cyp51, Nsdhl and Dhcr7*, and a reduced additive model *Age+Genotype+Sex* was used to fit data of fatty acid-‍related genes *Cyp7a1, Cyp8b1* and *Cyp27a1*. ANOVA type III test was used to estimate the statistical significance of the effects and Holm method was used to control FWER at α=0.05.

**Microarray-based gene expression profiling.** We hybridized 20 Affymetrix GeneChip™ Mouse Gene 2.0 ST Arrays (Affymetrix, Santa Clara, California, USA) with individual samples from livers of 14 mice (6 male, 8 female) at 24 months of age. 8 samples were obtained from WT mice, i.e. 4 from male and 4 from female mice. 6 samples were obtained from *Cyp51* KO mice, i.e. 2 from male and 4 from female mice. From each of *Cyp51* KO mice, two samples (a tumor and a surrounding tissue) were taken; altogether, 6 tumor and 6 corresponding surrounding-tissue samples were obtained from *Cyp51* KO mice (**Table ‍M3**).

#### Suppl. Table M3: Number of 24M mouse samples for DNA-microarray analysis. ↔ indicates related samples, i.e. different tissue samples obtained from individual mice.

| **24M mouse samples** | | **Genotype** | | | | |
| --- | --- | --- | --- | --- | --- | --- |
|  |  | *Cyp51* WT | *Cyp51* KO | | | |
|  |  | **Tissue** | | | | |
|  |  | Normal | Surrounding | | HCC | |
| **Sex** | Female | 4 | 4 | ↔ | | 4 |
|  | Male | 4 | 2 | ↔ | | 2 |

The experiment was performed with 500 ng starting concentration of total RNA with a purity (absorbance ratio A260/280) >1.8 and quality (RIN) >4.5. The procedure was done as described previously (2, 3) and considering the manufacturer’s instructions. Briefly, after overnight hybridization at 45 °C and 60 rpm/min microarrays were washed and stained using GeneChip Fluidics Station 450 and scanned on Affymetrix GeneChip Scanner 3000 7G. Image analysis and first quality check were done using the Affymetrix Expression Console™ version 1.3.

Further quality control and analysis were performed using R and Bioconductor software packages. RMA algorithm from package *oligo* (8) was used for the normalization of raw expression data. Package *arrayQualityMetrics* (9) was used to check additional quality control and assessment of potential outliers before and after normalization. Raw (CEL) as well as normalized data were deposited to GEO under accession number GSE127772 (<https://www.ncbi.nlm.nih.gov/geo/query/acc.cgi?acc=GSE127772>). Package *limma* (10) was used to fit individual normalized gene expression and gene set enrichment data using a linear regression model *Genotype+Sex+Genotype:Sex*. Empirical Bayes statistics was used to estimate the significance of the effects for each gene (gene set) and Benjamini-Hochberg procedure was used to control false discovery rate (FDR) at α=0.05 accounting for the number of tested genes (gene sets) and to infer differential expression of genes (enrichment of gene sets).

KEGG PATHWAY (10) and TRANSFAC database were used for functional enrichment studies (11). Gene sets containing over 5 elements were constructed and tested for enrichment using the *PGSEA* package (12). In the case of transcription factor enrichment, factors were merged based on their ID irrespective of their binding sites.

Gene expression data from 19 weeks (19W) old hepatocyte-specific *Cyp51* KO mice (*Cyp51^flox/flox^; Alb-Cre* or LKO; i.e. KO) and their WT littermates (*Cyp51^flox/flox^* or LWT; i.e. WT) following standard rodent diet (low-fat no-cholesterol, LFnC) was analysed to determine processes that contribute to tumor progression from 19W towards 24M. Data is available from GEO under accession number GSE58271 (<https://www.ncbi.nlm.nih.gov/geo/query/acc.cgi>) (2). Data was preprocessed, annotation updated and analyzed using the same procedure as described above. A linear regression model *Genotype+Sex+Genotype:Sex* was used to fit individual normalized gene expression and gene set enrichment data from mouse samples shown in **Table ‍M4**.

#### Suppl. Table M4: Number of 19W mouse samples from GSE58271 that were included to determine processes that contribute to tumor progression from 19W towards 24M.

| **19W mouse samples** | | **Genotype** | |
| --- | --- | --- | --- |
|  |  | WT | KO |
| **Sex** | Female | 3 | 3 |
|  | Male | 3 | 3 |

Network diagrams of differences in gene expression in females and males (**Fig. ‍4C**) were created using the NetworkAnalyst program (13). For the generation and visualization of transcription factor networks (**Suppl.** **Fig. 7**), the list of female and male DEG was mapped to the ˝database˝ libraries of the STRING online software (14) using medium confidence score (0.4).

***ADDITIONAL REFERENCES***

10.	Smyth GK. Linear models and empirical bayes methods for assessing differential expression in microarray experiments. Statistical applications in genetics and molecular biology. 2004;3:Article3. PubMed PMID: 16646809.

11.	Matys V, Kel-Margoulis OV, Fricke E, Liebich I, Land S, Barre-Dirrie A, et al. TRANSFAC and its module TRANSCompel: transcriptional gene regulation in eukaryotes. Nucleic acids research. 2006 Jan 1;34(Database issue):D108-10. PubMed PMID: 16381825. Pubmed Central PMCID: 1347505.

12.	Furge K, Dykema K. PGSEA: Parametric Gene Set Enrichment Analysis. R package version 1.44.0 ed ed.

13.	Xia J, Benner MJ, Hancock RE. NetworkAnalyst--integrative approaches for protein-protein interaction network analysis and visual exploration. Nucleic acids research. 2014 Jul;42(Web Server issue):W167-74. PubMed PMID: 24861621. Pubmed Central PMCID: 4086107.

14.	Szklarczyk D, Morris JH, Cook H, Kuhn M, Wyder S, Simonovic M, et al. The STRING database in 2017: quality-controlled protein-protein association networks, made broadly accessible. Nucleic acids research. 2017 Jan 4;45(D1):D362-D8. PubMed PMID: 27924014. Pubmed Central PMCID: 5210637.

15.	Long J, Wang H, Lang Z, Wang T, Long M, Wang B. Expression level of glutamine synthetase is increased in hepatocellular carcinoma and liver tissue with cirrhosis and chronic hepatitis B. Hepatol Int. 2011 Jun;5(2):698-706. PubMed PMID: 21484108. Pubmed Central PMCID: 3090553.

***SUPPLEMENTARY FIGURES***

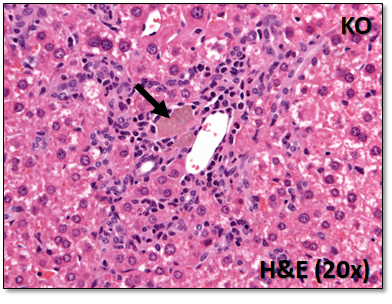

**Suppl. Fig. 1: Patho-histological features of *Cyp51* KO mice.** In livers of aging mice, yellow pigment likely representing lipofuscin pigment was observed in areas of pronounced ductular reaction, sometimes surrounded by other hepatic cells, evaluated as macrophages. Accumulation of yellow pigment in the liver of *Cyp51* KO mice (arrow), (H&E, original magnification x200); N=6-‍10 mice/group. KO, *Cyp51* KO.

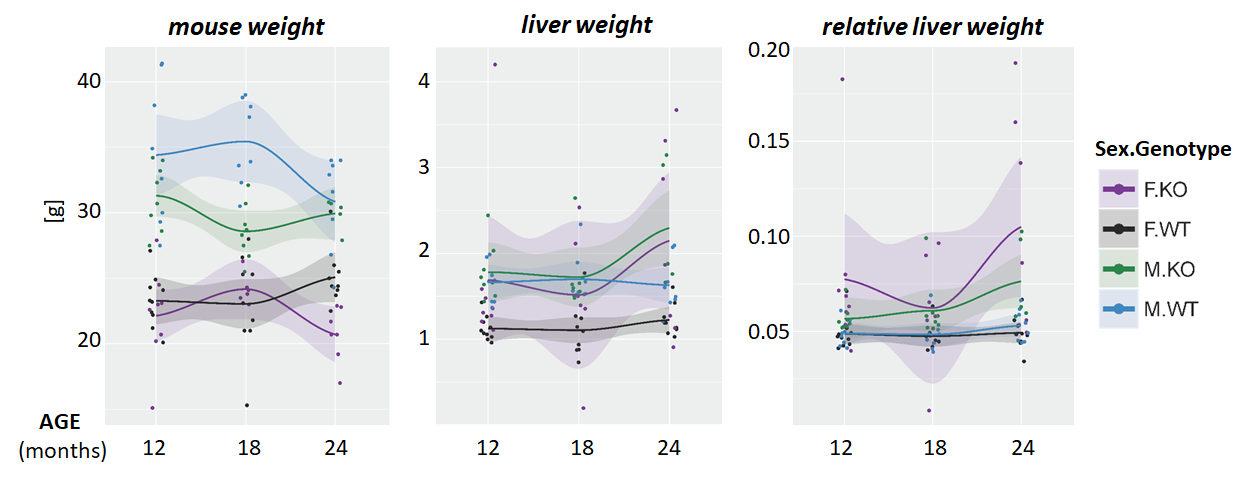

**Suppl. Fig. 2:** **Comparison of mouse liver/body/relative weights:** relative weights were significantly elevated in 24M old KO females, who had the highest incidence of liver tumors. The KO mice of both sexes have significantly increased liver and decreased body weight at different ages. 5-‍10 mice were analysed in each group. G, gram; KO, *Cyp51* KO; WT, *Cyp51* WT; F = female; M = male.

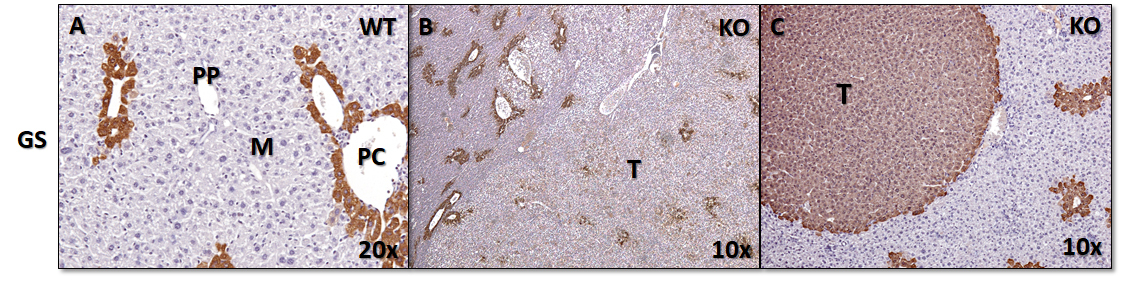

**Suppl. Fig. 3:** **Immunohistochemical expression of glutamine synthetase (GS)** as a recently proposed marker of HCC (15). As expected, GS staining was positive in pericentral hepatocytes (PC), but not in midzonal (M) or periportal (PP) hepatocytes in livers of *Cyp51* KO and *Cyp51* WT mice. In tumor tissue of *Cyp51* KO mice, different staining patterns were observed, from negative, focal, to diffuse strong staining.

The immunohistochemical reaction for GS in (**A**) WT mice showing staining in pericentral hepatocytes, while some tumors in KO mice showed weakly positive reaction (**B**) and some intensively positive reaction (**C**). GS, glutamine synthetase; T, tumor; PP, periportal; PC, pericentral, M, midzonal liver zone; KO, *Cyp51* KO; WT, *Cyp51* WT. Original magnification, **A** x200, **B** x100, **C** x400*.*

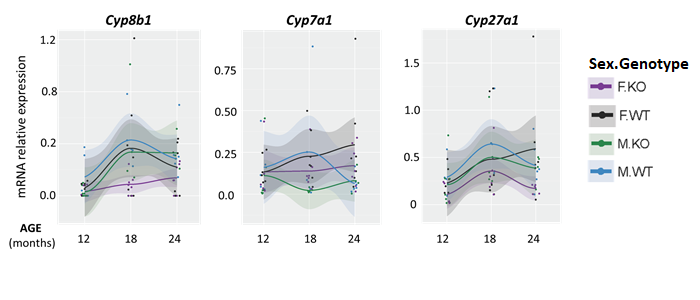

**Suppl. Fig. 4:** **Relative mRNA expression of selected genes** **of bile acid (BA) synthesis** represented different sex and age adaptation on *Cyp51* ablation.

Age profiles of qPCR expression measurements of genes selected for validation include BA synthesis genes (*Cyp8b1*, *Cyp7a1,* and *Cyp27a1*) (N= 5 per sex/genotype group) for different mice genotypes and sexes. Light color bands represent 95% confidence interval and dots represent individual measurements. KO, *Cyp51* KO; WT, *Cyp51* WT; F = female; M ‍= ‍male.

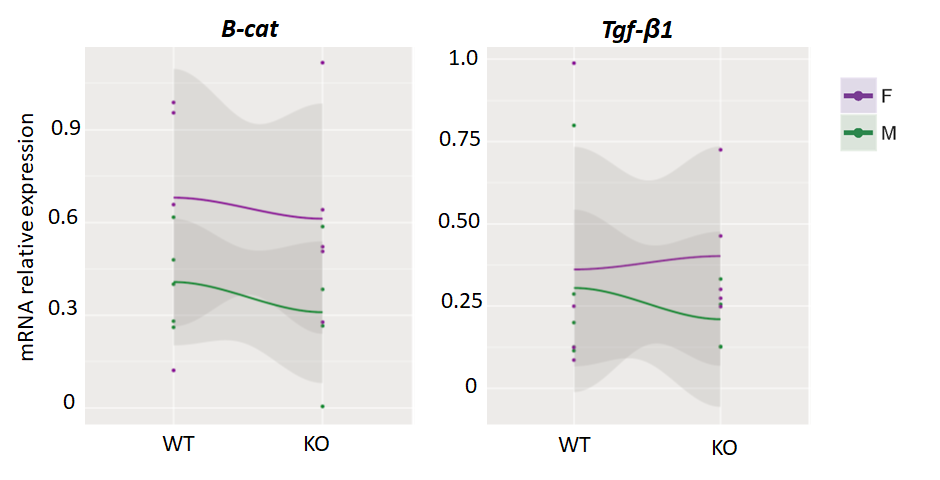

**Suppl. Fig. 5: qPCR expression profiles of TGF-β1 and Wnt signalling markers.** Genotype profiles of qPCR expression measurements of selected genes β-‍catenin and TGF-β1 in 24M KO mice. Light color bands represent 95% confidence interval and dots represent individual measurements. (N= 3-4 mice per sex/genotype group); KO, *Cyp51* KO; WT, *Cyp51* WT, β –‍cat, β-‍catenin; F, female; M, male.

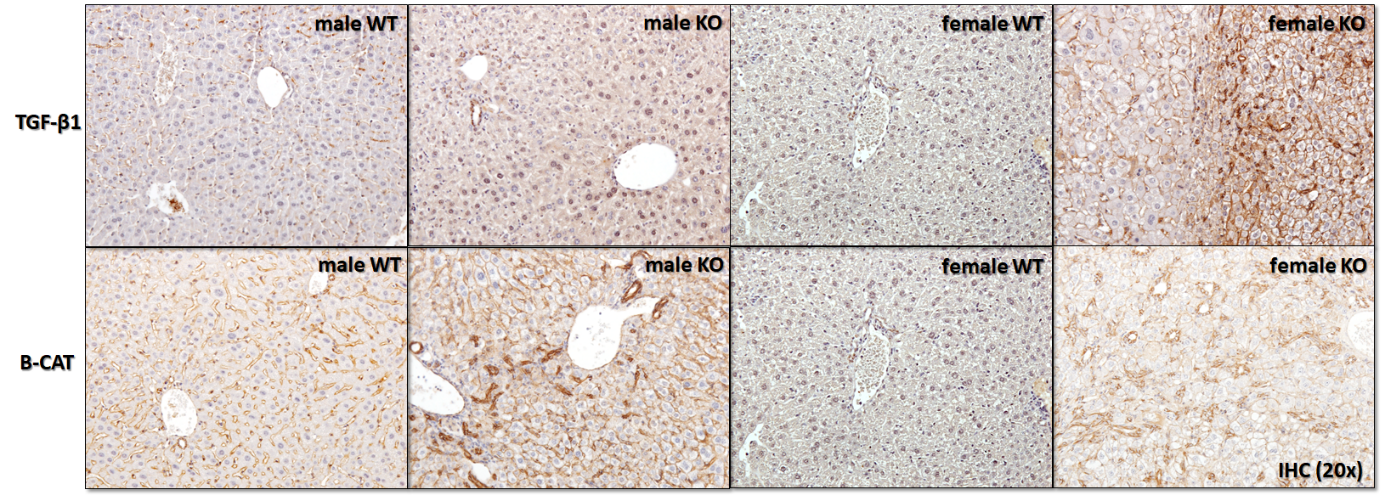

**Suppl. Fig. 6: Immunohistochemical expression of TGF-β1 and β-catenin** as potential markers of hepatocarcinogenesis in 24M.‍KO mice. A similar observation was made in 12M.KO mice (images not included). Original magnification, x200. β-CAT, β-catenin; KO, *Cyp51* KO; WT, *Cyp51* WT*.*

***24M KO enriched TF network***

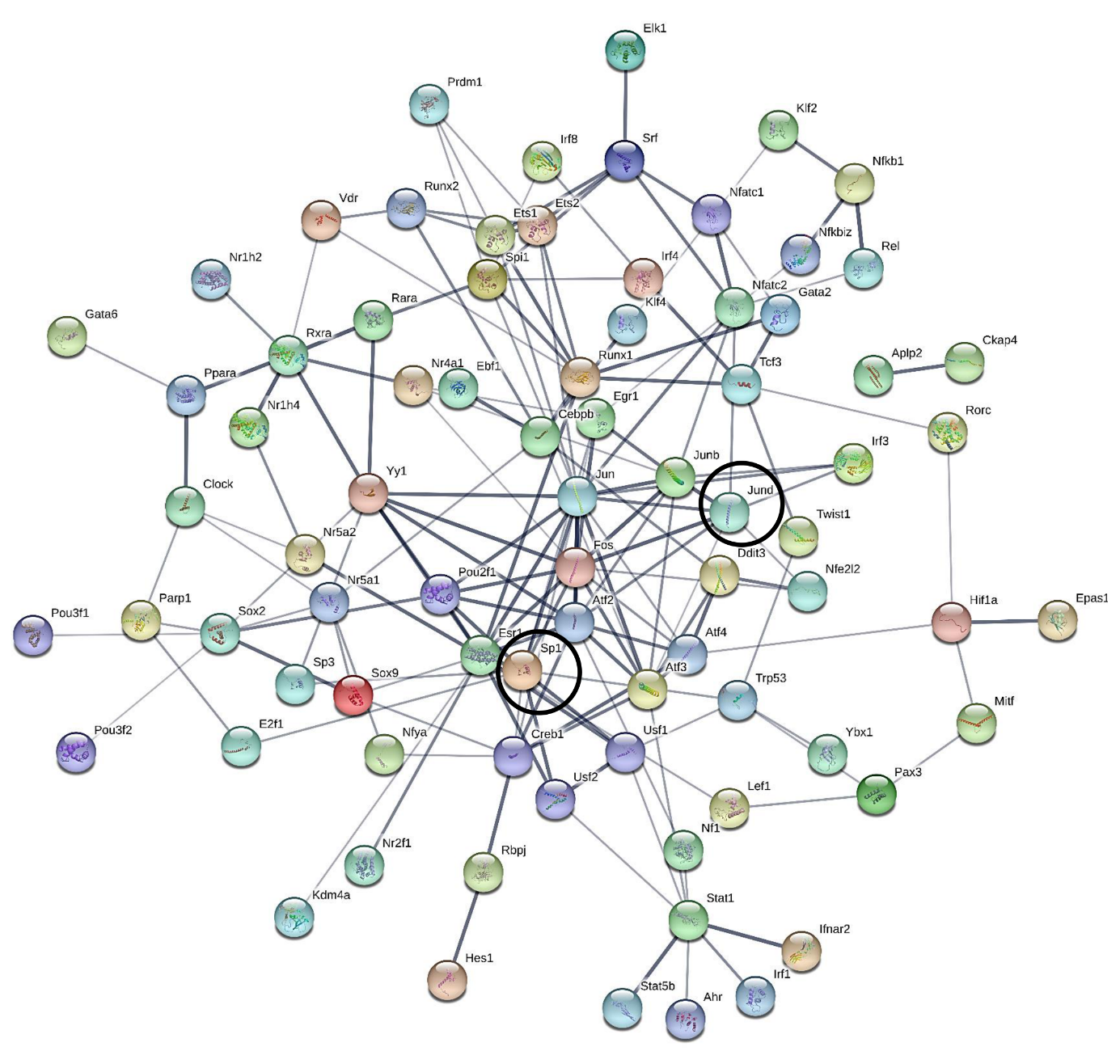

**Suppl. Fig. 7: TF networks of 24M.KO mice.**

TF networks generated with STRING show known and predicted protein-protein interactions. Network nodes represent TFs, produced by a single, protein-coding gene locus. For TFs with the known or predicted 3D structure, the structure is represented within nodes; empty nodes represent TFs of unknown 3D structure. Edges indicate protein-protein associations, i.e. proteins jointly contributing to a shared function and not necessarily physically binding each other. Nodules representing TFs SP1 and JUND are surrounded by black lines. KO, *Cyp51* KO.

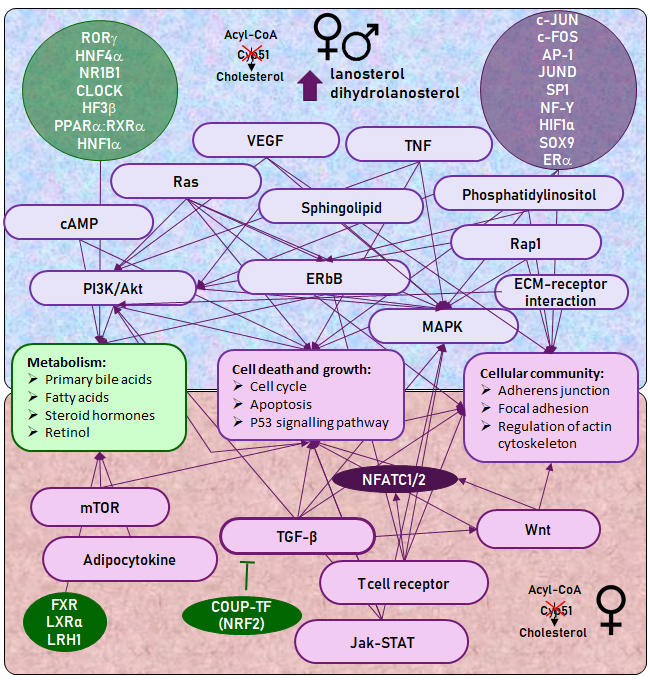

**Suppl. Fig. 8. Hepatocyte-specific *Cyp51* KO resulted in the deregulation of multiple signalling pathways and TFs leading to the development of female-‍prevalent liver cancer.** Arrows indicate connections between KEGG signalling pathways, TFs, and cellular processes. KEGG pathways and TFs are violet if positively enriched and green if negatively enriched at 24 months. **Top half** - selected KEGG signalling pathways and TFs that were deregulated in females and males, leading to inhibition of basic metabolism and increase in processes of cell death and growth, and cellular communication. MAPK and PI3K/Akt signalling pathways have a central role and are regulated by other signalling pathways and TFs, all directly or indirectly contributing to cancerogenesis. **Bottom half** - female-specific KEGG signalling pathways and TFs. TF NFATC1/2 is central to female-specific phenotype and is connected by TGF-β, mTOR and Wnt signalling pathways, positively enriched only in female *Cyp51* KO livers.
